# Great tits in noisy territories and avoiding overlapping respond stronger to territorial intruders

**DOI:** 10.1101/808733

**Authors:** Çağlar Akçay, Y. Kağan Porsuk, Alican Avşar, Dilan Çabuk, C. Can Bilgin

**Affiliations:** Department of Psychology, Koç University, Istanbul, Turkey; Department of Biological Sciences, Virginia Tech, Blacksburg, VA, USA; Department of Biological Sciences, Middle East Technical University, Ankara, Turkey

**Author notes:** corresponding author, Department of Psychology, Koc University, Rumelifeneri Kampusu, Sariyer 34450, Istanbul, Turkey. **Ethics:** All procedures in the study reported here were consistent with the ASAB/ABS guidelines for the treatment of animals in behavioural research and teaching. No subjects were captured or handled during the experiment and all procedures (including removing the flagging and noise measurements) ended within about 10 minutes of arriving at the territory to minimize disturbance. **Data, code and materials:** The data and R-script required to reproduce the results are available with the manuscript as supplementary files.

**Keywords:** song overlapping, urbanization, anthropogenic noise, aggressive signalling, threat signals

## Abstract

Animals often communicate with each other in noisy environments where interference from the ambient noise and other signallers may reduce the effectiveness of signals. Signalling behaviours may also evolve to interfere with signals of their opponents, e.g. by temporally overlapping them with their own, such as the song overlapping behaviour that is seen in some songbirds during aggressive interactions. Song overlapping has been proposed to be a signal of aggressive intent, but few studies directly examined the association between song overlapping and aggressive behaviours of the overlapping bird (the predictive criterion). In the present paper we examined the question of whether song overlapping is correlated with aggressive behaviours displayed during a simulated territorial intrusion in a population of great tits (*Parus major*) living in an urban-rural gradient. We also examined whether aggressive behaviours are correlated with the ambient noise levels. We found that overlapping was associated negatively with aggressive behaviours males displayed against a simulated intruder. These results fail to support the predictive criterion for song overlapping, raising the question whether overlapping is in fact a signal of aggressive intent. Ambient noise levels were associated positively with aggressive behaviours but did not correlate with song rate, song duration or song overlapping. Great tits in noisy urban habitats may display higher levels of aggressive behaviours due to either interference of noise in aggressive communication or another indirect effect of noise.

## Introduction

The environment through which signals are sent and received is open to interference by either sources of noise or other signals traveling through the medium (Bee and Micheyl, 2008; Brumm, 2006; Gil and Brumm, 2014; Wiley, 2006). Animals generally tend to space their signals temporally to minimize interference from other individuals or sources of noise (Brumm, 2006; Ficken et al., 1974; Gil et al., 2014; Wasserman, 1977; Wilson et al., 2016). In some circumstances however, temporally overlapping the signal of another individual may be a deliberate behaviour used as a signal. Indeed, overlapping a rival’s song has been proposed to be a signal of aggressive intent used in territorial defence by songbirds (Dabelsteen et al., 1997; Mennill and Ratcliffe, 2004; Naguib and Mennill, 2010).

Signals of aggressive intent (also called threat signals) are signals that are not costly to produce but carry information about the likelihood of escalation by the signaller (Caryl, 1979; Searcy et al., 2006). In a recent review, Searcy and Beecher (2009) laid out three empirical criteria for a signal to be established a signal of aggressive intent: 1) the signal should occur more often in agonistic interactions than other non-agonistic interactions (context criterion); 2) it should correlate with other aggressive behaviours or predict a subsequent escalation (predictive criterion) and 3) the presence or absence of it should elicit different responses in receivers (response criterion).

For the most part, research on song overlapping as an aggressive signal focused on the response criterion, asking whether birds respond differently when their songs are being overlapped (vs. not overlapped) experimentally via interactive playback, i.e. the response criterion (see reviews in Helfer and Osiejuk, 2015; Naguib and Mennill, 2010; Searcy and Beecher, 2009). Generally, these studies do find changes in responses of subjects when their songs are being overlapped (vs. not overlapped) that are consistent with the hypothesis that subjects perceive overlapping as an aggressive challenge (Naguib and Mennill, 2010).

Fewer studies addressed the question whether song overlapping is correlated with aggressive behaviours or subsequent escalation, i.e. the predictive criterion (see table S1 for more details of these studies). We focus on this latter set of studies, because our main aim is to assess the predictive criterion. Among these studies, only two found a positive correlation between song overlapping measures and approach measures that are generally taken as indicators of aggression (Brindley, 1991; van Dongen, 2006). Three studies found no correlation between song overlapping and approach (Baker et al., 2012; Fitzsimmons et al., 2008; Yang et al., 2014). Finally, two studies found a negative correlation between song overlapping and some approach measures (Vehrencamp et al., 2007; Wilson et al., 2016). Thus, on balance of it, it is not clear whether the predictive criterion for song overlapping is satisfied among the various species that has been studied to date (Table S1).

A second aim of our study is to ask whether interference due to ambient noise also has an effect on aggressive behaviours. Although noise is a feature of natural habitats as well, the impact of noise in animal communication has garnered special attention due to the increasing levels of anthropogenic noise that affects social behaviour of wildlife (Gil and Brumm, 2014; Johnson and Munshi-South, 2017; Shannon et al., 2016). Anthropogenic noise may make signals less detectable (Kleist et al., 2016; Templeton et al., 2016) or less effective (Halfwerk et al., 2011). In response, animals may change the frequency, amplitude or length characteristics of their signals (Brumm and Todt, 2002; Slabbekoorn and den Boer-Visser, 2006; Wood and Yezerinac, 2006) or change their signalling or social behaviours (Halfwerk et al., 2012; Halfwerk and Slabbekoorn, 2009).

If noise interferes with effective signalling during aggressive interactions than we may expect signallers to display higher levels of aggression in noisier habitats. Consistent with this, in several species of songbirds birds living in noisier urban habitats display higher levels of aggression than birds living in rural habitats (Davies and Sewall, 2016; Evans et al., 2010; Fokidis et al., 2011; Foltz et al., 2015; Hardman and Dalesman, 2018). Recent studies also found more direct evidence for a positive correlation between ambient noise and aggressive responses of territory holders (Grabarczyk and Gill, 2019; Phillips and Derryberry, 2018; Wolfenden et al., 2019), although at least one study found a negative correlation between noise and aggressive responses (Kleist et al., 2016).

In the present study, we aim to assess the predictive criterion for song overlapping in great tits *(Parus major).* While song overlapping has long been proposed to be a signal of aggressive intent in great tits (Amy et al., 2010; Dabelsteen et al., 1996; Langemann et al., 2000; Peake et al., 2001), to the best of our knowledge no previous study has explicitly examined whether this behaviour is correlated with aggressive behaviours of the overlapping bird in this species. Some studies (Amy et al., 2010; Snijders et al., 2017; Snijders et al., 2015) did measure approach behaviours and song overlapping of the same males via playback experiments but these studies did not report analyses that would directly pertain to the predictive criterion. We also assess whether ambient noise is correlated with aggressive responses during territorial defense. Finally, we ask (in supplementary materials) whether song overlapping, song rate, and song duration also varies with ambient noise.

## Methods

### Study site and subjects

We studied 42 male great tits holding territories on the campus of Middle East Technical University, in suburban Ankara, Turkey (39°53’32” N, 32°47’03” E). Although the campus is located in a steppe habitat, it includes a large area that has been afforested mostly with conifers in the last six decades. Great tits nest in human structures on campus (e.g. access ports left open in telephone and electricity poles or cavities on the side of buildings) as well as nest boxes provided in the forested parts and natural cavities spread throughout the campus grounds. We located the territories of great tits by observation of singing posts during song recording (see below) or locating nests in the boxes or human structures. For birds whose territories we determined from singing posts, we observed each bird for 15-30 minutes and occasionally used playbacks to determine the extent of their territory (these observations were not done on the day of trial but 2-3 days prior). For territories with known nests (n=17, most of them being in nest boxes but some, n= 6, in other human-made cavities; none had nestlings) we carried out playbacks within 5m of the nest. The birds were not captured at any time and therefore were not banded. Each bird was tested once. The trials were carried out between 7 and 13 April 2019 at the start of the nesting period in the morning hours between 0600 and 1130. We avoided testing neighbouring males on the same day and carried out consecutive trials in locations that are more than 300 m apart (where the previous subject was out of earshot for the experimenters).

### Playback Stimuli

We recorded male songs from great tits on the research site in Spring 2018 and 2019 using a Marantz PMD660 or 661 recorder and a Sennheiser ME66/K6 shotgun microphone. Using the software Syrinx (John Burt, Seattle, WA) we viewed and selected high quality recordings of songs to create playback stimuli. After manually filtering out low-frequency noise (–<~>1000 Hz), we added a silent period to create a 7-second wave file that was played in loop such that playback rate was one song every 7 seconds. The average (±SD) playback song duration was 2.99 (±0.42) seconds (range= 2.01-3.55 seconds, corresponding to an average duty cycle of 0.42). We created 24 playback stimuli (see Figure S1 in supplementary materials for an example stimulus). For each subject we used a stimulus song that was recorded at least 1km away from the subject’s territory which is considered a stranger song in previous studies on great tits (Falls et al., 1982). This constraint meant we used some stimulus songs multiple times for different males (11 stimuli were used in two trials, 2 stimuli were used in three trials, and one stimulus was used in 4 trials-the rest of the songs were used once).

### Procedure

We placed a wireless speaker (Anker SoundCore, Anker, Inc) face-up at a natural perch at about 1.5-2m from the ground, inside the territory of the male (either 5 m from the nest if the nest was known or at a location that was central to the various singing posts the male was observed to be singing).

We then set up flags at 1m, 3m and 5m distance on either side of the speaker to help with distance estimation during the trial. The observers then stepped back to about 15 m from the speaker. The playbacks were controlled from a smartphone connected to the speaker via Bluetooth.

The trials were recorded with the same equipment as above. We started the trials by playing back the stimulus song until the subject responded either by singing a song or approaching the playback (the subject was within 15-20 m from the speaker in the first response). After this first response, we continued the playback for another 3 minutes and we narrated the behaviour of the subject by noting each flight, distance to the speaker and singing behaviours. The behaviours during this 3-minute period are the main response variables of the study.

### Noise measurements

After the trial, we removed the flagging, took a GPS reading (Garmin eTrek, Garmin Inc.) of the trial location and carried out the ambient noise measurements using the method described by Brumm (2004). Briefly, we took two measurements in each cardinal direction at the location of the playback using a sound level meter (VLIKE VL6708, VLIKE Inc.) with A weighting and fast response (125 msec) settings for a total of 8 measurements that lasted between 1-2 minutes. We averaged these measurements to get an average noise level. In a subset of territories (n=32 out of 42) we took a second measurement of ambient noise later in the day (between 1300 and 1600) to assess how consistent variation in the ambient noise was. For logistical reasons we could not take second noise measurements in 10 territories. The first and second noise measurements were highly correlated with each other (linear regression: β=0.94, SE= 0.17, t=5.46, p= 6.4 × 10^-6^; intra-class correlation coefficient: r=0.64, p< 0.00001). We used the first noise measurement in the analyses as this measurement was available for all territories.

### Response measures

Viewing the trial recordings in Syrinx, we extracted the following behaviours: flights (all airborne movements), distance to speaker with each flight, and songs. Because the trial durations slightly varied across trials (ranging between 179 seconds to 192 seconds), we converted the counts of flights and songs into rates by dividing it with the trial duration. We also calculated the proportion of the trial subjects spent within 1 m of the speaker and noted the closest approach of the distance.

Definition of aggressive behaviors is a significant issue in studying signals of aggressive intent (Searcy and Beecher, 2009). This issue was also recognized by Naguib and Mennill (2010) in their forum article on song overlapping (e.g. on page e2). We further note that in defining aggressive behaviors for the purpose of assessing whether a putative signal carries information on aggressive intent, it is necessary that these aggressive behaviors are defined separately from that putative aggressive signal so as to avoid a circular logic (Akçay et al., 2015; Searcy and Beecher, 2009). In the following, we define signals in the same way Otte (1974) did: “ behavioral, physiological, or morphological characteristics fashioned or maintained by natural selection because they convey information to other organisms” (p.738). Song and aspects of song (such as timing, rate, overlapping etc.) are unambiguously signals, as they have no function other than potentially conveying information to receivers. We define non-signaling aggressive behaviors as approaching the opponent, staying in close proximity and flying around (presumably in search of the opponent) or over the opponent, in line with previous research on territorial songbirds, including our focal species (e.g. Hardman and Dalesman, 2018; McGlothlin et al., 2007; Rivera-Gutierrez et al., 2010; Wingfield, 1994). These behaviors have been found to predict physical attack on dummies in several studies with songbirds (Akçay et al., 2014; Akçay et al., 2013; Araya-Ajoy and Dingemanse, 2014; Searcy et al., 2006; Searcy and Beecher, 2009). Therefore, we take the spatial behaviours (flight rates, closest approach distance and time spent within 1 m of the speaker) as indicators of aggressive response in the present paper.

For each song that the subject sang, we determined whether the song overlapped the playback song and calculated the proportion of the subject’s songs that overlapped the playback song. This measure therefore corrects for variation in song rate as suggested by Searcy and Beecher (2009), but it is undefined for subjects that did not sing any songs (n=7). Because of variation in duty cycle (duration of stimulus with song/entire duration of stimulus) between stimulus songs, we also classified subjects based on whether the observed song overlapping (as defined above) was higher (n=11) or lower (n=24) than the duty cycle (stimulus song duration/7 seconds). We classified the subjects who did not sing any songs as a third category in this variable. Note that the categorical song overlapping variable (over or under duty cycle) represents categorization with respect to a null model given by the duty cycle, while absolute levels of overlapping (proportion of subject’s song that overlapped the playback) gives the amount of overlapping relative to a perfect turn-taking case (i.e. zero overlapping). Thus, these two variables represent observed overlapping levels relative to two distinct “ null” models. We remain agnostic with respect to the appropriate null model, particularly because both analyses show the same results (see Results). Finally, for each subject we measured duration of each of their song and calculated the average song duration for each subject (see supplementary materials).

### Data Analysis

The rates of flights, closest approach distance and proportion of time spent within 1m of speaker were all highly correlated with each other (Table 1). We therefore used a principle component analysis (PCA) using package *psych* in R (Revelle, 2015) to arrive at a single measure of aggressive response strength. The first component of the PCA (PCA1, eigenvalue= 2.07) explained 68.95% of variation and was taken as the measure of aggressive response strength (hereafter called aggression score for brevity). The first component was positively correlated with flight rates and time spent within 1m, and negatively correlated with closest approach distance (Table 1).

**Table 1:** Pearson correlation coefficients (p-values) for the correlation between the aggressive response variables and their loadings to the first component of principle component analysis (PCA). Kaiser-Meyer-Olkin measure and Barttlet’s test were significant (KMO: 0.661; Bartlett’s test: χ^2^=33.86; p<0.00001)

|  | **Flight rate** | **Proportion of trial spent within 1m** | **Closest approach** |
| --- | --- | --- | --- |
| **Flight rate** | — | 0.60 ($2 \times 10^{-5}$) | -0.57 ($7 \times 10^{-5}$) |
| **Proportion of trial spent within 1m** | — | — | -0.42 (0.005) |
| **Loading Coefficient** | 0.88 | 0.81 | -0.79 |

To assess the relationship between song overlapping and aggressive responses, we ran a linear mixed model (LMM) with aggression score (PCA1) as the response variable, stimulus song as a random factor (to control for the multiple presentation of some stimuli), and song overlapping, defined as proportion of subject songs that overlapped the playback song as the fixed factor. This model had a reduced sample size as song overlapping defined as above was not defined for 7 subjects that sang no songs leaving 35 subjects. We then ran a second LMM on aggression scores with stimulus as a random factor and the following fixed factors: the categorical overlapping variable (with three categories: overlapping higher or lower than the duty cycle or no song) and ambient noise levels. This model included all subjects (n= 42). Finally, we ran a third LMM on aggression scores with the stimulus as random factor and song rate as fixed factor (we analysed song rates separately as our categorical overlapping variable had a no-song category that confounded with song rate). Further analyses on song variables and ambient noise are reported in the supplementary materials. All analyses were carried out in R using the package nlme (Pinheiro et al., 2017; R Core Team, 2012). We supply the R script as well as the data necessary to recreate the analyses reported.

## Results

Song overlapping (proportion of subject songs that overlapped the playback songs) was negatively related to the aggressive behaviours: high overlapping was indicative of lower levels of aggression scores (LMM: coefficient ± SE = −1.29 ± 0.52, χ2=6.27, p=0.012; Figure 1a). In the model with the categorical overlapping variable (Table 2, Figure 1b), both overlapping and ambient noise significantly predicted aggressive behaviours: birds that overlapped the playback less than expected had higher aggression scores compared to high-overlapping birds (unpaired t test: t(33)=2.78, p=0.0088). Birds who did not sing any songs in turn had higher aggression scores than low-overlapping birds (unpaired t-test: t(29)=2.10, p=0.044. Figure 1b).

**Figure 1.**
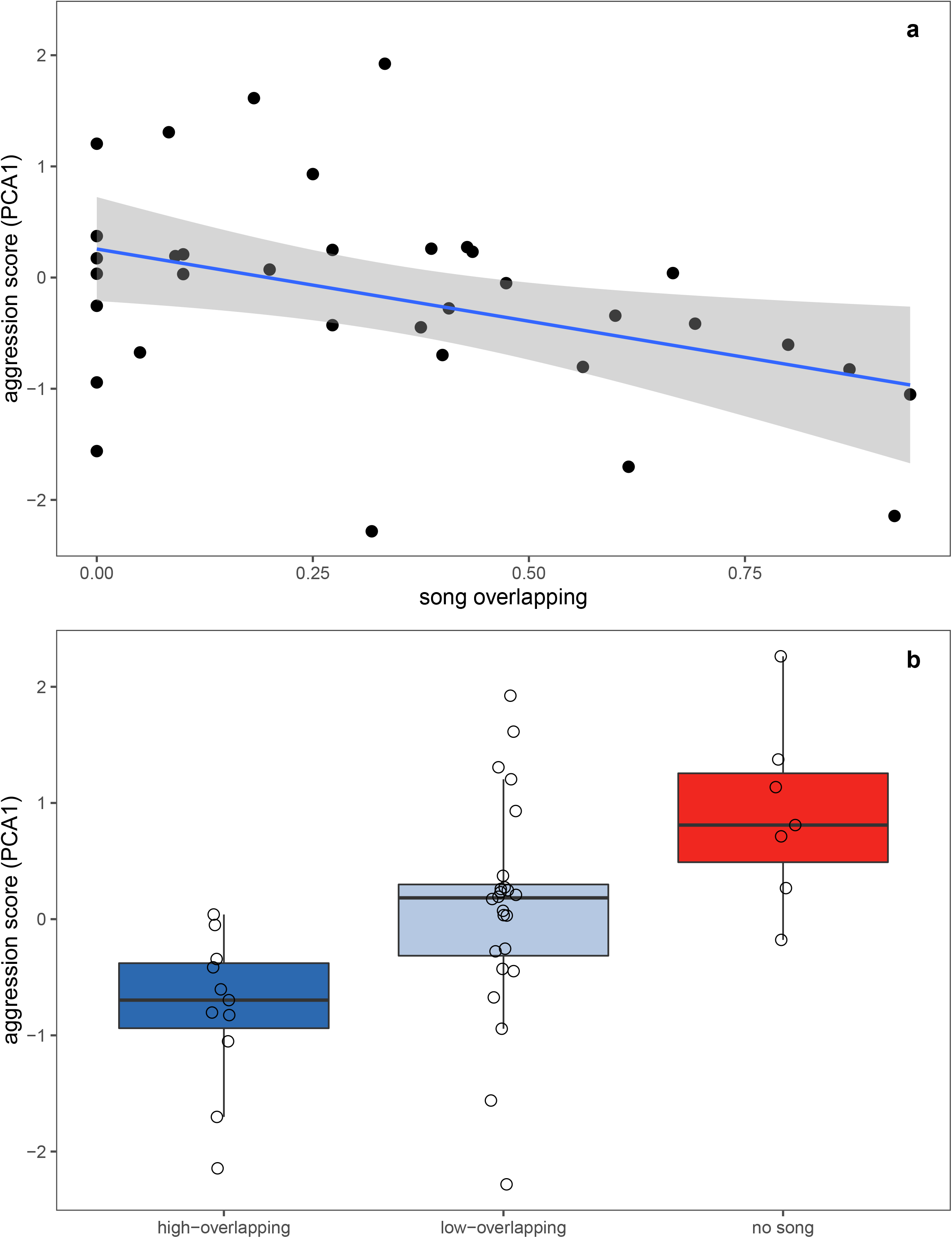
a) Scatterplot of song overlapping (proportion of the subject’s songs that started while the playback song was playing), and aggression scores (PCA 1). b) Aggression scores of subjects overlapping the playback songs at more than the duty cycle (n=11), less than the duty cycle (n=24), singing no song (n=7). The boxes indicate interquartile ranges and median, the whiskers indicate 95% confidence intervals and the open circles are the data for individual subjects.

**Table 2.** Linear mixed model (LMM) with stimulus song as random factor and aggression scores as response variable (n= 42 subjects). The predictor variables were song overlapping as the fixed factor (higher than duty cycle, lower than duty cycle or no-song; with higher than duty cycle taken as baseline) and ambient noise levels.

| **Predictor variable** | **Level** | **Coefficient (SE)** | **$\chi^2$** | **p-value** |
| --- | --- | --- | --- | --- |
| **Overlap (baseline= high)** | low | 0.65 (0.31) | 16.41 | 0.00027 |
|  | no song | 1.58 (0.39) |  |  |
| **Ambient noise** |  | 0.053 (0.022) | 6.01 | 0.014 |

Looking at ambient noise variation and aggressive responses, we found that birds living in noisier territories had higher aggression scores (Table 2, Figure 2a). The effect of song rate was significant and negative: birds that sang less displayed higher levels of aggressive behaviours (LMM: coefficient ±SE: 0.11 ± 0.048, χ^2^= 5.58, p=0.018; Figure 2b). Ambient noise was not correlated with any singing behaviours (Table S2 in supplementary materials).

**Figure 2.**
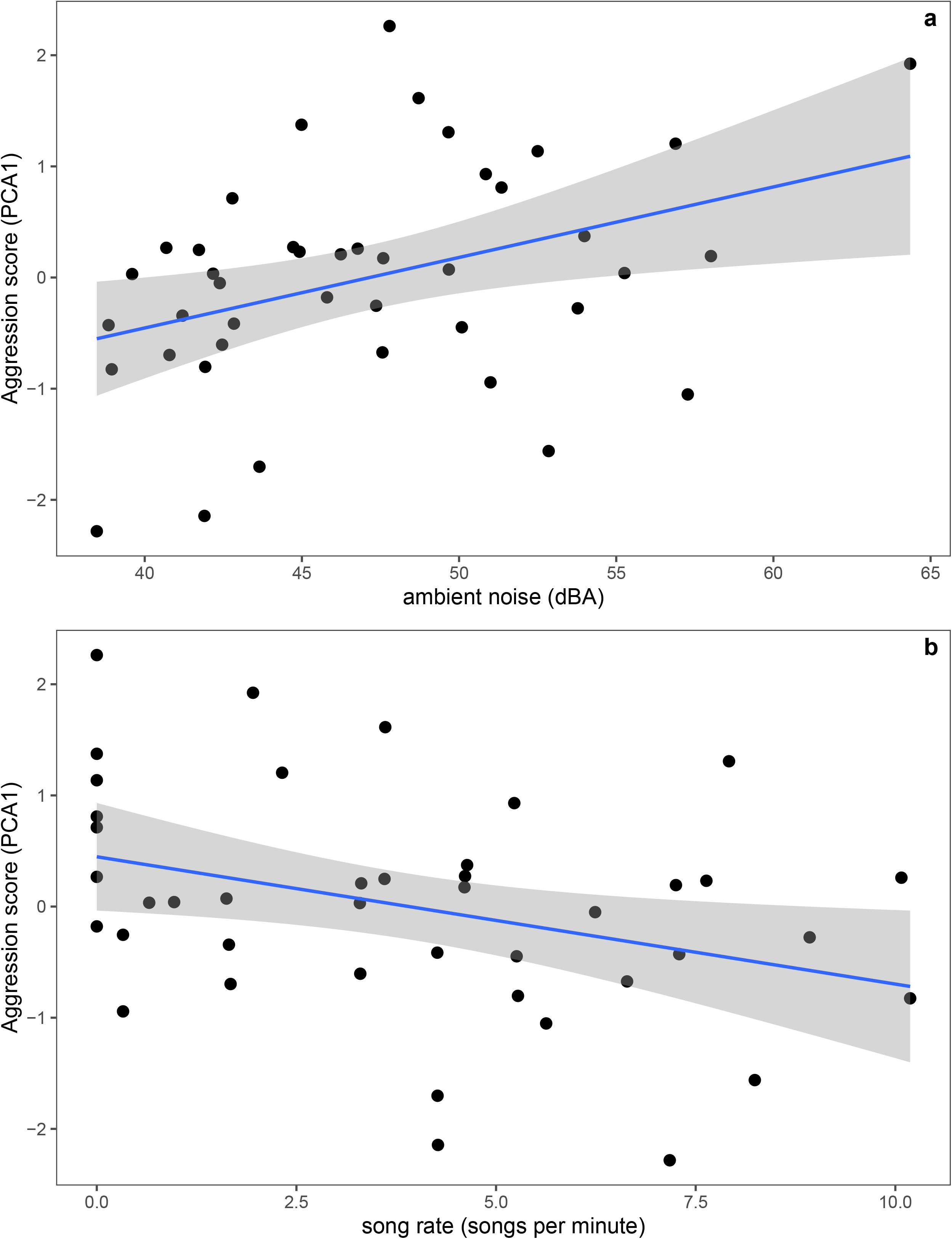
a) Scatterplot of ambient noise levels (average of 8 measurements with A settings, dB) and aggression scores (PCA1). b) Scatterplot of song rates and aggression scores (PCA1)

## Discussion

We set out to assess the predictive criterion for signals of aggressive intent as laid out by Search and Beecher (2009). This criterion suggests that a signal of aggressive intent should positively correlate with aggressive behaviours; our finding of a negative correlation between aggressive behaviours and song overlapping does not conform to this prediction.

### Is song overlapping an aggressive signal in great tits?

One limitation of our study is the relatively short playback period (3 minutes) and the absence of a physical model that the subjects could attack, we can’t rule out the possibility that song overlapping is an aggressive signal, but only in the early stages of escalation. Under this hypothesis, subjects that overlapped the playback song were simply not yet escalating to a high level of aggression by approaching the speaker, whereas the subjects that have approached the speaker closely were already at a high level of escalation. This possibility of overlapping being a slower escalation signal needs empirical testing, ideally in an escalating playback design (Akçay et al., 2013; Searcy et al., 2013). Nevertheless, we have some preliminary evidence that suggests this is unlikely: in a separate experiment involving five minutes of playback and 3D printed models of a great tit, aggression scores measured in the same PCA method as in the present study were predictive of an eventual attack on the model (Akçay, unpublished data). Thus, the aggression scores we report here seem to be reliable indicators of an impending attack.

Thus, the present study adds to the body of equivocal evidence on the predictive criterion for song overlapping reviewed in the introduction (Table S1) and to the best of our knowledge it is the first study to test the predictive criterion for song overlapping explicitly in great tits. Several previous studies collected both measures of aggressive behaviors (e.g. approach) along with overlapping but they did not report analyses that would test the predictive criterion directly. Amy and colleagues (2010) measured responses of male great tits (in terms of both approach and song overlapping, among other measures) to overlapping and alternating song playbacks but they do not report any analyses correlating song overlapping levels with aggressive responses. Similarly, Snijders and colleagues report two experiments (Snijders et al., 2017; Snijders et al., 2015) where song overlapping and approach behaviours data are collected but not analysed together to determine possible associations between these behaviours. Instead the focus on these studies have been on correlations of vocal behaviours (including song overlapping) and approach behaviours with exploration scores (see also; Jacobs et al., 2014).

In summary, previous studies in great tits, while often measuring song overlapping and approach measures together, did not explicitly present evidence that song overlapping indeed predicts or correlated with approach, even though many explicitly assume song overlapping to be an aggressive signal (e.g. Amy et al., 2010, p. 3686). Indeed, in a recent synthesis, Snijders and Naguib (2017) wrote that “ frequent signal overlap has been shown most often, yet not always, to reflect more aggressive intentions” (p. 319). In light of the equivocal evidence on predictive criterion in general (reviewed above and in Table S1), as well as the current findings in great tits specifically, we caution that this conclusion may be premature.

### Alternative hypotheses on song overlapping

Other hypotheses can explain both the differential responses to being overlapped (the evidence related to response criterion) and the equivocal evidence on the association between aggressive behaviours and overlapping (the predictive criterion). One possibility is the interference avoidance hypothesis (Wilson et al., 2016). A recent study on black-capped chickadees *(Poecile atricapillus)* found that males were less likely to overlap playback of songs when they were close to the speaker and more likely to overlap quieter stimuli compared to louder stimuli (Wilson et al., 2016). Interestingly, chickadees avoided overlapping both biotic (chickadee song) and abiotic (white noise) stimuli. Wilson and colleagues (2016) interpret these results as compelling evidence that overlapping in chickadees being not a signalling behaviour but a strategy to avoid interference with the signal (see also Baker et al., 2012). They then reinterpret the differential singing responses to being overlapped found earlier in this species, i.e. evidence for the response criterion (Mennill and Ratcliffe, 2004) as birds attempting to avoid interference from the overlapping signal (see Wilson et al., 2016 for a fuller discussion). Given the present results that present a similar pattern in great tits, we believe it would be prudent to consider non-signalling hypotheses for this species as well.

Another possibility is that overlapping is a signal but indicates another trait of the signaller, such as some aspect of genetic quality, personality or condition (Helfer and Osiejuk, 2015). For instance, the study by Snijders et al. (2017) found that males that were older and in better condition had higher vocal response scores (in which number of overlapping songs was a loading variable). Note however that their measure confounded song rate and overlapping and therefore it is not clear whether song overlapping independent of song rate varies with condition or age. Interestingly, another study on great tits found that birds that were exposed to an ectoparasite as nestlings were found to be less likely to overlap the song playback carried out in the territory after they were recruited to the population (Bischoff et al., 2009). Song overlapping however, was not correlated with the condition of the male as a nestling, and there was only a trend towards a positive association between overlap and male condition as an adult. Thus, while this study suggests a developmental constraint, the mechanism for this constraint is unclear.

Overlapping could also serve as a signal of personality variation among males (Amy et al., 2010; Snijders et al., 2015). Several studies looked at correlations between overlapping and personality variation in great tits measured via exploration scores in a novel environment test. Snijders et al. (2015) found their vocal response measure (a PCA that includes number of overlaps and song rates, and therefore does not control for song rate) was positively correlated with exploration scores. Amy et al. (2010) found that the correlation between exploration scores and proportion of playback songs that were overlapped by the subject (which again confounds song rate) was positively or negatively associated with exploration score depending on whether the playback was trying to overlap the subject or not. Finally, Jacobs et al. (2014), studying another population of great tits, found that exploration scores was not related to proportion of playback songs that subjects overlapped. In this last experiment birds with slower exploration scores displayed higher levels of aggressive behaviours, unlike in the Amy et al. (2010) study which found the opposite effect. All in all, the results on personality variation and overlapping present no clear picture.

Because we did not have information on our subjects’ age or condition, we could not test any hypothesis that might relate song overlapping to individual traits other than aggressive intent. More data are clearly needed to test these possibilities. For instance, the hypothesis that song overlapping is a signal of individual quality implies that individuals should display repeatable song overlapping when they are challenged multiple times, but as far as we are aware this possibility has not been tested.

### Noise and aggressive behaviours

A second aim of our study was to assess the effect of noise on aggressive behaviours. We found a positive correlation between aggressive behaviours and ambient, consistent with the hypothesis that noise may play a significant role in determining aggressive phenotypes observed in urban populations of birds (Akçay et al., 2019; Davies and Sewall, 2016; Evans et al., 2010; Hardman and Dalesman, 2018). The effect of noise on aggression may come about via multiple mechanisms. First, noise may disrupt effective communication (McMullen et al., 2014; Ríos-Chelén et al., 2015) which in turn may increase aggressive behaviours (Logue et al., 2010).

Interestingly, unlike other species (Brumm and Slater, 2006; Ríos-Chelén et al., 2013) great tits in our population did not show any evidence that they change their song rate, average song duration or overlapping depending on ambient noise levels. It is worth noting that urban noise tends to be in the lower frequency ranges (less than 2kHz); whereas great tit songs tend to have minimum frequencies above 2 kHz (see Figures S1 and S2). It is therefore possible that the lack of an effect of urban noise on song rates, duration and overlapping may be due to a relatively low masking due the noise. Previous research however showed conclusively that songs in great tits and other songbirds become less effective in urban noise (Halfwerk et al., 2011; Luther et al., 2015). Furthermore, great tits were found to switch to song with higher minimum frequencies when they were experimentally presented with low frequency urban noise suggesting that they actively try to avoid masking from the noise (Halfwerk and Slabbekoorn, 2009). Noise also have a general effect on making animals increase the amplitude of their signals, termed the Lombard effect (Brumm and Slabbekoorn, 2005) and urban birds in several species including great tits are found to sing songs with higher minimum frequencies (Mockford and Marshall, 2009; Slabbekoorn and Peet, 2003; Wood and Yezerinac, 2006). Whether the latter phenomenon comes about due individual plasticity, developmental plasticity or longer term selection of higher frequency songs is an area of active research (Moseley et al., 2018; Zollinger et al., 2017). Whatever the mechanism, these findings suggest that acoustic communication is likely hampered in noisier territories which may result in higher aggression.

At the mechanistic level, noise may also lead to higher levels of aggression via increased physiological arousal or stress, as has been documented in humans (Baron and Richardson, 2004). In particular, exposure to noise may elevate baseline levels of glucocorticoids (Creel et al., 2002; Crino et al., 2011; Davies et al., 2017; but see Kleist et al., 2018; Zollinger et al., 2019) which in turn may lead to a higher level of aggression in individuals living in noisier areas. Interestingly, in a recent study on urban and rural song sparrows, Davies and colleagues (2018) found that baseline levels of corticosterone (the main avian glucocorticoid) in blood samples after a simulated territorial intrusion did not differ between urban and rural birds, and the levels of corticosterone were positively related to territorial aggression males displayed. For a given level of plasma corticosterone however, urban birds displayed more aggression than rural birds, suggesting that urban birds potentially show differences in glucocorticoid signalling from rural birds. Thus, the exact physiological mechanisms that determines the relationship between urbanization and noise on the one hand, and aggression on the other is yet unclear (Bonier, 2012). Further experimental studies are needed to quantify the effect of noise on plasticity at behavioural and physiological levels to tests hypotheses on the physiological mechanisms of aggression and how signalling strategies change under noisy conditions.

### Conclusion

In summary we found that song overlapping in great tits did not satisfy the predictive criterion for being an aggressive signal. Our findings on ambient noise meanwhile, corroborate the recent studies linking noise and aggression and suggest new avenues to investigate the effect of anthropogenic noise on animal social behaviour and the physiological mechanisms underlying it. Together, the current study highlights how animals may manage interference from noise or other signallers in their social interactions.

## Acknowledgements

We would like to thank Mike Beecher, Bill Searcy and Marc Naguib for comments on the manuscript.

