## Supplementary Materials for "Great tits in noisy territories and avoiding overlapping respond stronger to territorial intruders"

Table S1: Studies that explicitly test the predictive criterion for song overlapping in territorial songbirds. With the exception of Fitzsimmons et al. which analyzed natural singing interactions with an acoustic location system, all studies used simulated intrusions via playback. Van Dongen (2006) used a caged live decoy along with playback. Baker et al. used a taxidermic mount with playback. The other studies used only playbacks.

| Study | Species | Measure of Song Overlap | Measure of aggressive response | Finding |
|---|---|---|---|---|
| **Brindley, 1991[1]** | European robins | "Proportion of total songs given by subject bird found to overlap those of either neighbor" | PCA (including closest approach, time spent close, latencies to sing and to approach, and number of songs sang during and after) | Birds respond more strongly to strangers than neighbors, and also overlap strangers more than neighbors. |
| **van Dongen, 2006[2]** | golden whistler | Whether the subject song overlapped the playback song or not | Distance to the simulated intruder | Birds were closer to the simulated intruder when they were overlapping than when they were not. |
| **Baker et al. [3]** | Black-capped chickadees | Proportion of subject songs that overlapped the playback song | Attack on a taxidermic mount | Attackers did not have higher rates of overlapping than non-attackers either during the entire trial or in the 1 minute before attack. |
| **Fitzsimmons et al.[4]** | Black-capped chickadees | Whether two birds overlapped each other during natural interactions | Approach towards each other | "overlapping itself was not related to subsequent approach in escalated contests" (p.1918) |
| **Yang et al. [5]** | Eurasian wrens | Proportion of subject songs that overlapped the playback song | Closest approach distance, latency to sing after the playback | "Neither the responders' closest approach distances to the speaker nor their latencies to sing after the playback started were correlated with their overlapping levels" (p.87) |
| **Vehrencamp et al. (2007)[6]** | Banded wrens | Proportion of subject songs that overlapped the playback song | Latency to first approach, latency to retreat, closest approach, proportion of time spent within 15m | High overlapping was associated with early retreat and not associated with other approach variables |
| **Wilson et al. (2016)[7]** | Black-capped chickadees | Whether or not the subject overlapped the playback song | Distance to the speaker | Subjects were farther away from the speaker when they overlapped the playback song |

*Comparison of duty cycles and observed overlapping levels.*

We compared the duty cycle of the stimulus song each subject received with their observed overlapping levels to provide a test of whether song overlapping was observed more or less frequently than expected by chance. We recognize that different "null" models will lead to different results [8] and our aim is not to be comprehensively test and show that song overlapping happens more or less expected by chance. Rather we present this analysis for the sake of completeness. The main conclusion in the paper (that individual variation in song overlapping does not correlate with aggressive behaviors) does not depend on any particular choice of null model.

We compared the duty cycle of stimulus to observed overlapping with a paired t-test. The observed levels of overlapping (mean ± SD: 0.33 ± 0.29) was lower than the duty cycle of stimulus songs (0.42 ± 0.06) although not significantly so; paired t-test: $t(34)=-1.89$, $p=0.068$.

*Analyses on ambient noise levels and signing behaviors*

Here we report analyses on ambient noise levels and song overlapping, song rate, and average song duration. Several studies found that noise has an effect on singing behaviours, most notably in change of amplitude and frequency [9-11], but also song rate [12] and song duration [13]. Song overlapping may also change as a result of ambient noise: individuals may be less likely to overlap their opponent's songs in noisy environments if ambient noise already interferes with the transmission of the signals. Alternatively, noise may make it hard to reliably detect start and end of opponent signals [14], which may increase overlapping rates. To the best of our knowledge, no previous study examined whether song overlapping varied with ambient noise levels.

To assess the question of whether ambient noise was associated with singing behaviors we ran seperate LMMs on song rate, song overlapping and average song duration as response variables, and ambient noise levels as predictor variables, with stimulus song as a random factor, to determine whether ambient noise had an effect on other singing behaviours. The model on average song duration and song overlapping again had a reduced sample size (n= 35) because these variables were not defined for subjects that did not sing any songs.

The results showed that ambient noise was not related to song overlapping, song rate or average song duration (Table S2).

Table S2. Linear mixed models on song rate, average song duration and song overlapping as response variables and ambient noise as predictor variable. Stimulus song is a random factor in all models.

| ssModel | $\chi^2$ | p-value |
|---|---|---|
| **Song rate** | 0.063 | 0.80 |
| **Song duration** | 0.69 | 0.79 |
| **Song overlapping** | 2.18 | 0.13 |

Figure S1. Sonogram and amplitude spectra of a great tit song. Note that the lower frequencies (<1000 Hz) were filtered manually in Syrinx. Sonogram produced with “rspect" function in R by Matt Wilkins, PhD. Amplitude spectra was produced with function meanspec in the package seewave in R.

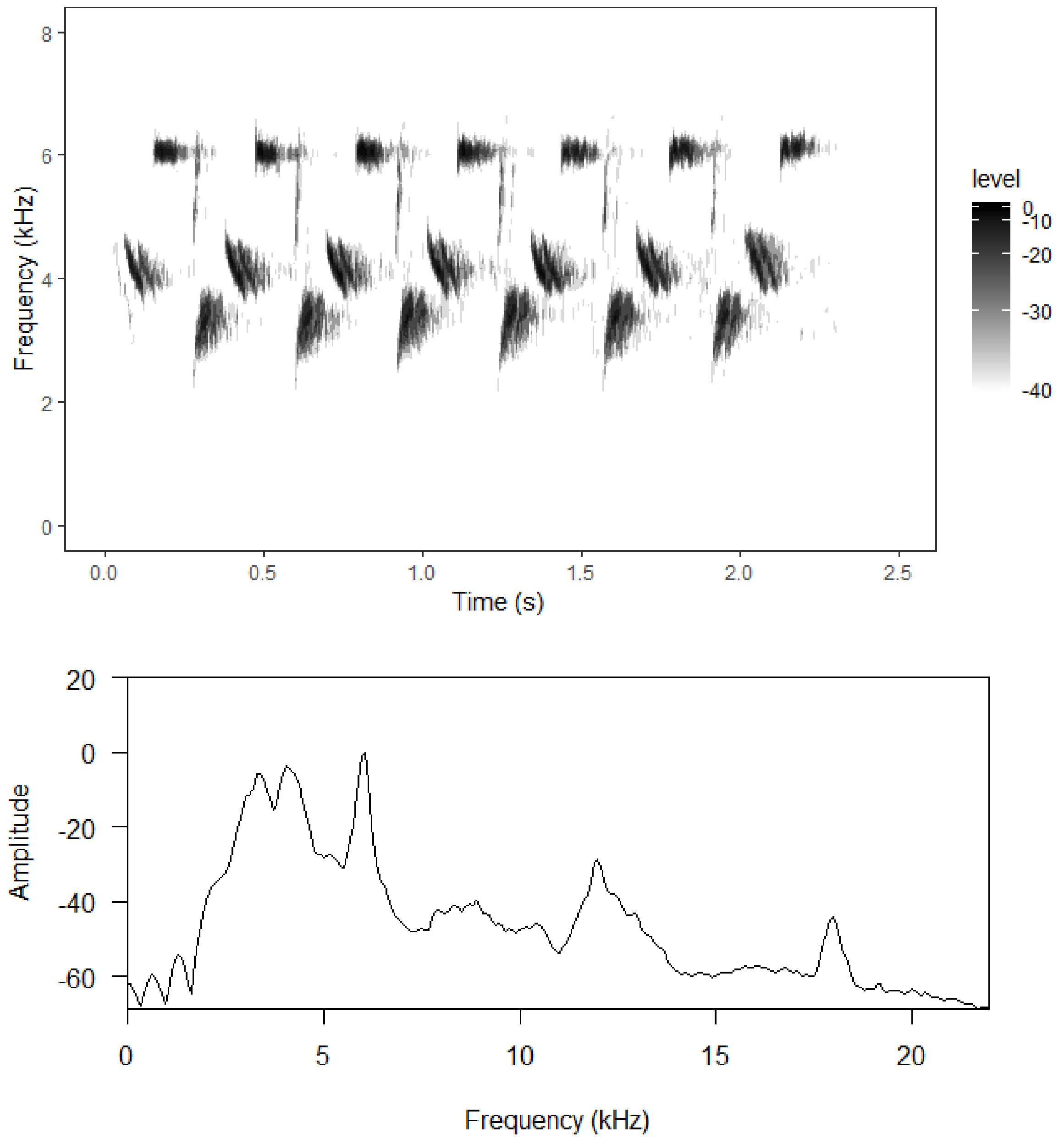

Figure S2. The amplitude spectrum of background noise in an urban territory.

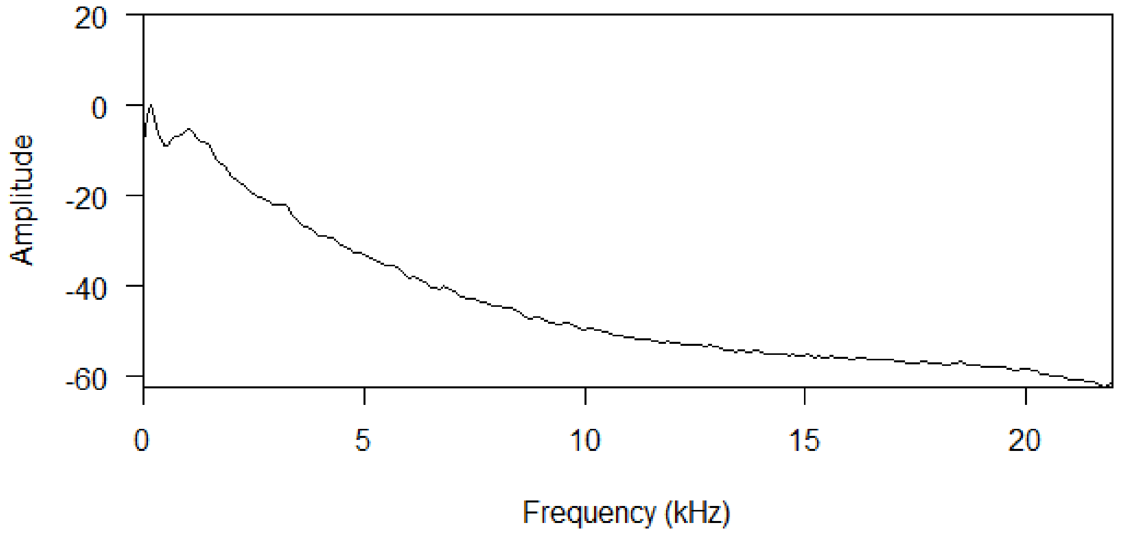
